# The stoichiometric interaction model for mesoscopic molecular dynamics simulations of liquid-liquid phase separation

**DOI:** 10.1101/2022.04.19.488761

**Authors:** Yutaka Murata, Toru Niina, Shoji Takada

## Abstract

Liquid-liquid phase separation (LLPS) has received considerable attention in recent years for explaining the formation of cellular biomolecular condensates. The fluidity and the complexity of their components make molecular simulation approaches indispensable for gaining structural insights. Domain-resolution mesoscopic model simulations have been explored for case in which condensates are formed by multivalent proteins with tandem domains. One problem with this approach is that interdomain pairwise interactions cannot regulate the valency of the binding domains. To overcome this problem, we propose a new potential, the stoichiometric interaction (SI) potential. First, we verified that the SI potential maintained the valency of the interacting domains for the test systems. We then examined a well-studied LLPS model system containing tandem repeats of SH3 domains and proline-rich motifs. We found that the SI potential alone cannot reproduce the phase diagram of LLPS quantitatively. We had to combine the SI and a pairwise interaction; the former and the latter represent the specific and non-specific interactions, respectively. Biomolecular condensates with the mixed SI and pairwise interaction exhibited fluidity, whereas those with the pairwise interaction alone showed no detectable diffusion. We also compared the phase diagrams of the systems containing different numbers of tandem domains with those obtained from the experiments, and found quantitative agreement in all but one case.

**SIGNIFICANCE:** Cells organize their interior structures as not only membrane-bound organelles but also as membrane-less organelles. Membrane-less organelles, such as stress granules, Cajal bodies, and postsynaptic density, are biomolecular condensates in which many biomolecules are gathered owing to their interactions. In some cases, biomolecular condensates are formed by tandemly connected multidomain proteins. In such cases, a mesoscopic simulation model representing each domain as a particle is effective; however, the problem with this approach is that a domain-domain pairwise interaction cannot regulate the well-defined valency. To overcome this problem, in this study, we have developed a new potential, viz. the stoichiometric interaction potential, and confirmed that this potential can reproduce the liquid-liquid phase separation of multidomain proteins, a hallmark of the membrane-less organelles.

## INTRODUCTION

In the last decade, biomolecular condensates have gained considerable attention in a broad range of cellular biology-related areas (1). Biomolecular condensates are non-stoichiometric large assemblies of biomolecules, and are most often composed of proteins and RNAs (2). Biomolecular condensates include membrane-less organelles known for a long time, such as Cajal bodies, nuclear speckles, stress granules, and postsynaptic density (3–6). More recently, biomolecular condensates have been suggested in many other contexts, such as signal transduction and chromatin organization (7). Many condensates exhibit liquid-like properties and are formed via the liquid-liquid phase separation (LLPS). Although not easy to be strictly proven, plausible functions of formation of these condensates have been suggested and examined. The formation of condensates leads to an increase in the local concentration of the involved proteins, which facilitates reactions of these proteins. The LLPS efficiently segregates many proteins into those inside (high-density phase) and outside (low-density phase) the condensates, enabling selective reactions. Recent studies have suggested a role of buffering; the concentrations of proteins in the low-density phase are maintained at a nearly constant level, thus increasing the robustness of the environment(8). Phase transition can lead to a sharp increase in the signal transduction response.

Biomolecular condensates are formed via a network of intermolecular interactions. A class of condensates is formed by intrinsically disordered proteins with promiscuous interactions throughout the protein, such as electrostatic, π-π, cation-π and hydrophobic interactions(9). In this case, the elementary interaction is not stoichiometric. In another class, proteins possess multiple globular domains connected by linkers, and each domain interacts with domains/modules of other proteins/RNAs(10). Molecular networks in the postsynaptic density are examples of this case(11). This class uses domain-domain stoichiometric interactions, which are most often one-to-one, interactions. Multi-valency per protein plays a key role in this case(12).

Since most of interactions in biomolecular condensates are fragile and dynamic, it is very challenging to characterize them experimentally. Molecular dynamics (MD) simulations are useful tools to compensate for this problem. Therefore, numerous simulation models and methods have been proposed in recent years. First, given the number of proteins and timescales involved, conventional all-atom MD simulations are currently beyond the reach. Coarse-grained protein models in which each amino acid is represented by one particle have been successfully applied to simulate LLPS and condensates made by the promiscuous interaction of intrinsically disordered proteins (Dignon et al.(13), Das et al.(14), Lafage et al.(15), Chatterjee et al.(16), Tesei et al.(17))(18).

On the other hand, for biomolecular condensates formed by stoichiometric interactions of globular domains, the above residue-resolution coarse-grained models are rather time consuming and thus less effective. Instead, theoretical modeling and mesoscopic model simulation have been more successful (8, 19–23). However, the theoretical model does not fully capture the flexible nature of the linkers as well as the spatial correlation among domains. Mesoscopic models represent each globular domain as a particle. Among mesoscopic models, the lattice and the patch-particle models are concise ways to realize the stoichiometric interactions. In the lattice model, stoichiometric interaction is automatically realized by the lattice (19, 24–26). However, the lattice representation has some limitations in representing a variety of domain sizes and the interacting networks. In the patch-particle model, the stoichiometric interaction is realized by adding a small patch of attraction (27–29). The introduction of the patch, however, largely reduces the time step. The reaction-diffusion framework also provides useful mesoscopic simulation models (30). The reaction-diffusion model, while useful, requires a simulation framework that is significantly different from standard MD simulators.

In this study, to simulate biomolecular condensates via domain-wise stoichiometric interactions, we propose a new stoichiometric interaction model for mesoscopic modeling of biomolecules. The stoichiometric interaction model takes a simple function, so that we can rely on standard MD methods. The stoichiometric interaction takes a function similar to that described by Lu et al. (31), as well as that of the Generalized-Born implicit solvent model(32, 33). This is not a pairwise additive, but its computational cost is same to that of pairwise interaction. Using an experimentally well-characterized binary mixture of tandem repeats of SH3 domains and the proline-rich modules (PRM), we validated the stoichiometric interaction model. We examined the liquid nature of the droplet, simulated the spontaneous assembly of the condensates. We also quantified the LLPS phase diagrams with several different stoichiometries and found quantitative agreement in all but one case.

## THEORY AND METHODS

### Meso-scale Model

To simulate a mixture of multidomain proteins, we, herein, introduce a mesoscopic model in which each domain of proteins is represented by one spherical particle. For example, SH3_4_, a tandem repeat protein with four SH3 domains, was modeled as four particles connected by three linkers. The linker was modeled using a linker potential *V*_*linker*_. The domains of different proteins interact via non-bonded interactions, which were decomposed into a pairwise term *V*_*pair*_ and a non-pairwise term called the stoichiometric interaction *V*_*SI*_. Thus, the total potential energy function was generally written as,

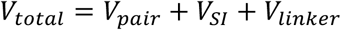

In the comparative studies mentioned below, we sometimes turned off one of the first two terms. The detail of each term is explained below.

For time propagation, we used a simple underdamped Langevin dynamics realizing the *NVT* ensemble where the temperature and friction coefficient are 300 K and 1.62 × 10^−4^ τ^-1^ respectively. In this case, τ ∼ 49 fs is base time unit of the system where energy is kcal/mol, length is Å, and mass is AMU. Note, however, that this time scale is used merely in the simulation software and does not mean the physical time. The mapping to actual time can be done by matching the diffusion coefficient from the simulation and that from the Stokes-Einstein formula for the SH3 domain and water viscosity. This resulted in τ∼128.5ps. We used the BAOAB Langevin integrator (34, 35).

### Pairwise Interaction

*V*_*pair*_ is a pairwise interaction applied to all the pairs of particles in a system. This is represented by the following equation:

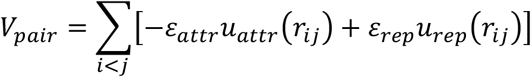

Here, the attractive potential is represented by a truncated cubic term

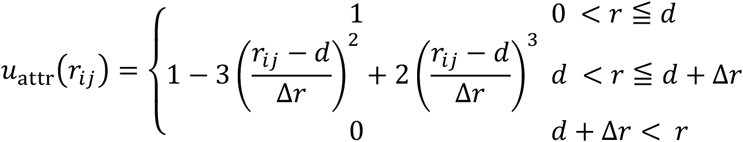

where *r*_*ij*_ is the distance between the two particles *i* and *j, d* is the sum of the radii of the two particles, and Δr is the range of the applied attractive force. We used 10.0Å as Δr for all simulations.

The short-range repulsive potential is defined by the Weeks-Chandler-Andersen (WCA) potential

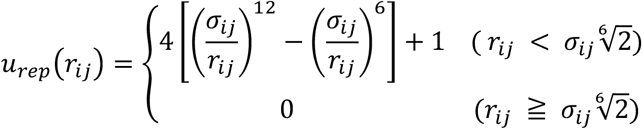

where σ_ij_ is calculated by

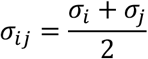

σ_i_ is twice the radius of gyration (Rg) of the domain; the actual values were SH3: 0.994nm and PRM: 0.896nm. The Rg of the SH3 domain was calculated from its structure (PDBID:4WCI). Rg of the PRM motif was calculated by a preliminary coarse-grained MD simulation. In this preliminary simulation, each amino-acid was represented by one particle. The potential function included a simple local potential (the flexible local potential, FLP)(36, 37) and non-local potential representing the excluded volume and electrostatics. Electrostatics was modeled by the Debye– Hückel theory.

We turned off the attraction between the same type of particles setting ε_attr_ as 0.0 (kcal/mol) throughout this study. For different type particles, i.e., between SH3 and PRM, ε_attr_ was set to 3.91 (kcal/mol) for the pairwise-only simulation and to 0.5 (kcal/mol) for the simulation with the Stochiometric Interaction Potential. These values were tuned to reproduce the experimentally determined binding affinity between SH3 and PRM (the procedure will be described later). We set ε_rep_ to 0.6 (kcal/mol) for all particle pairs.

### Worm-Like Chain Potential

*V*_*linker*_ was used to model a flexible linker using Worm-like chain model.

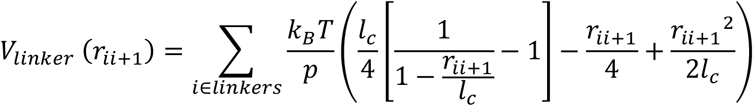

where k_B_ is the Boltzmann constant, *T* is the temperature, *r*_*ii*+1_ is the distance between particles i and i+1 connected by a linker, *p* is the persistent length, and *l*_*c*_ is the maximum length of the linker polymer. We set *p* equal to 0.39 (nm) and *l*_*c*_ as 0.38 (nm) based on a previous report (38).

### Stoichiometric Interaction Potential

In this study, we propose a stoichiometric interaction (SI) potential that modulates the valency of the domain-domain interactions. The SI is defined as

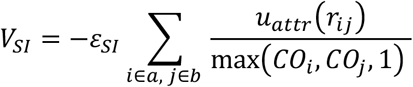

where the i-th particle is of type a, and the j-th particle is of type b. Here, a and b represent the types of domains. In the present study, these were either SH3 or PRM. *CO*_*k*_ is the contact number of the k-th particle and is defined for the one-to-one valency case as follows:

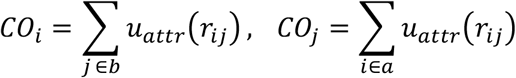

For more general n-to-m valency case, *CO*_*k*_ is defined by

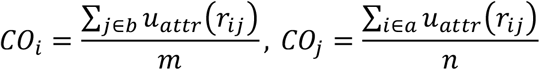

We set ε_SI_ to 3.91 (kcal/mol) throughout this study. The max() is a simple max function used in this study. The function *u*_*attr*_ () is the same as that of pairwise interaction with Δr = 10.0Å. The choice of Δr value has arbitrariness; we chose a reasonable value and checked the robustness of the results with respect to this value (Fig. S1).

While a naive implementation of the SI potential leads to a computational cost with an order higher than pairwise interaction, we can avoid it by carefully writing a code, as described in the Supporting Information. We also note that the simple max function is not differentiable, which causes inaccuracy in numerical simulations with a large time step. While we used this nondifferentiable form in this work with use of a modest time step, the simple max function can be replaced with the soft-max function when one wants to avoid this issue.

### Interaction Coefficient Tuning

The interaction strength parameter ε_attr_ (or ε_SI_) between SH3 and PRM was tuned to fit previously reported dissociation constant *K*_*d*_ = 356 *µM* reported previously (39). We considered that one SH3 domain was placed at the center of a sphere and one PRM was distributed around it. The radius of the sphere was determined so that the SH3 (and PRM) concentration inside the sphere was 712 μM. Under this condition, the contact probability between SH3 and PRM should be 1/2 to match the dissociation constant. Thus, we could tune the value of ε_attr_ by numerically solving the following equation,

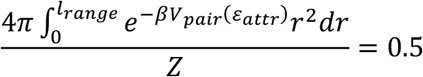

where *l*_*range*_ denotes the contact threshold, 1.95 nm. *β* = 1/(*k*_*B*_*T*) where *T* is 300 K. Z is the partition function defined as follows:

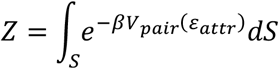

where S is the sphere around the SH3 particle. Using this procedure, we obtained the resulting value ε_attr_ (or ε_SI_) = 3.91 (kcal/mol).

### Stoichiometry Assay

We examined the stoichiometry realized by the pairwise and SI potentials using a molecular system containing 1000 SH3 and 1000 PRM monomers in a cubic box. The box size was set such that the concentrations of SH3 and PRM were 4mM. We tested three cases of attractions: 1) only the pairwise interaction, 2) only SI potential, and 3) no attractive interaction (as a control). In all cases, the WCA repulsive interactions were applied. Starting from a random configuration, Langevin MD simulations were performed with a time step of Δt=2.0. Each run consisted of 10^7^ MD steps. We repeated the same simulation five times with different stochastic forces. For the analysis, we used every 10^4^ steps in the last 10^6^ steps of the entire trajectory. The contact threshold was 1.95 nm. For the case of valency being 1:2 and 2:3 case, we just performed simulations with the only SI potential case.

### Direct Coexistence Method

To analyze the phase diagram of SH3-PRM systems, we used the direct coexistence (DC) method (27). In the DC method, we prepared a phase-separated configuration in a slab box, as follows: first, to build a condensed phase configuration, we placed SH3_n_ and PRM_n_ molecules, with the total number of domains being 3000, into a 25.0 × 25.0 × 25.0 nm^3^ box, which was surrounded by a repulsive wall potential. Given this, we ran 10^6^ step MD simulations for equilibration with Δt=2.0. Then, we moved the final configuration into a slab box of 25.0 × 25.0 × 415.0 nm^3^ with a periodic boundary condition. Finally, we performed the production runs of 10^8^ steps with Δt=2.0 were performed. All the simulations were conducted using an in-house program named Mjolnir, which is freely available on GitHub (https://github.com/Mjolnir-MD/Mjolnir).

### Phase concentration calculation

We divided the slab box into 50 bins perpendicular to the long axis and calculated the concentration in each bin. The dilute phase concentration was the mean of concentrations of which bins have concentrations lower than the mean of all bin concentrations. The condensed phase concentration was the mean of concentrations of all other bins.

### Condensed phase detection

To determine the presence and the size of the condensed phase at each step, we first built a contact matrix between SH3 and PRM domains. For this matrix, the threshold for the domain contact was 1.95 nm. From the domain-wise contact matrix, we obtained the molecule-wise contact matrix and listed up the moleculer clusters. The size of the largest cluster was defined as the size of the condensed phase. The condensed phase in MSD Analysis was defined as this largest cluster.

### MSD Analysis

For the mean square displacement (MSD) calculations, we set up a system that included 3000 particles (375 molecules) of SH3_4_ and PRM_4_ and ran a 10^8^ step MD simulation with Δt=5.0 using the DC method. Notably, molecules could move between the condensed and low-density phases during the simulation. To distinguish the MSD in the two phases, we needed to carefully pick up the test molecules. We divided the last 5.0×10^7^ steps into 10^5^ step time windows without overlap. At the beginning of each time window, we selected the SH3 and PRM molecules that were closest to the center of mass of the condensed phase and calculated the MSD for these SH3 and PRM. For the dilute phase, we calculated the MSD for those SH3 and the PRM, which were the farthest from the center of mass of the condensed phase. In the cases of the dilute phase, we removed 13 of 500 windows from the calculation because the target particle moved from the dilute phase to the condensed phase. In other case, there is no transition between the condensed and dilute phases. For the diffusion coefficient calculation, we calculated MSD for all steps in the window and fitted this curve for the range from 31 to 100 steps by open-source Julia package LsqFit.jl (https://github.com/JuliaNLSolvers/LsqFit.jl).

## RESULTS AND DISCUSSION

### Pairwise interaction cannot represent stoichiometry of protein-protein interaction

We chose an experimentally well-characterized two-component system composed of SRC homology 3 (SH3) domains and proline-rich motif (PRM) ligands as a model system for LLPS (39, 40). These domains often appear as tandem modules in signaling proteins. First, we focused on the SH3_4_-PRM_4_ system, where both SH3_4_ and PRM_4_ are composed of four domains connected in tandem. Previous studies have shown that the critical concentration for the LLPS is between 100-200μM for the SH3_4_-PRM_4_ system (39).

In our mesoscopic model, each domain was represented as one sphere, and the neighboring domains were connected by a linker potential (Fig. 1A) (see Methods for details). For analyzing the interaction between the SH3 and PRM domains, we first employed a simple pairwise interaction in which the attraction interaction was tuned to reproduce the experimentally determined dissociation constant *K*_*d*_ = 356 μM (Fig. 1B)(39). We simulated a phase-separated system using a slab box containing 375 molecules of both SH3_4_ and PRM_4_ (Fig. 1C).

**FIGURE 1.**
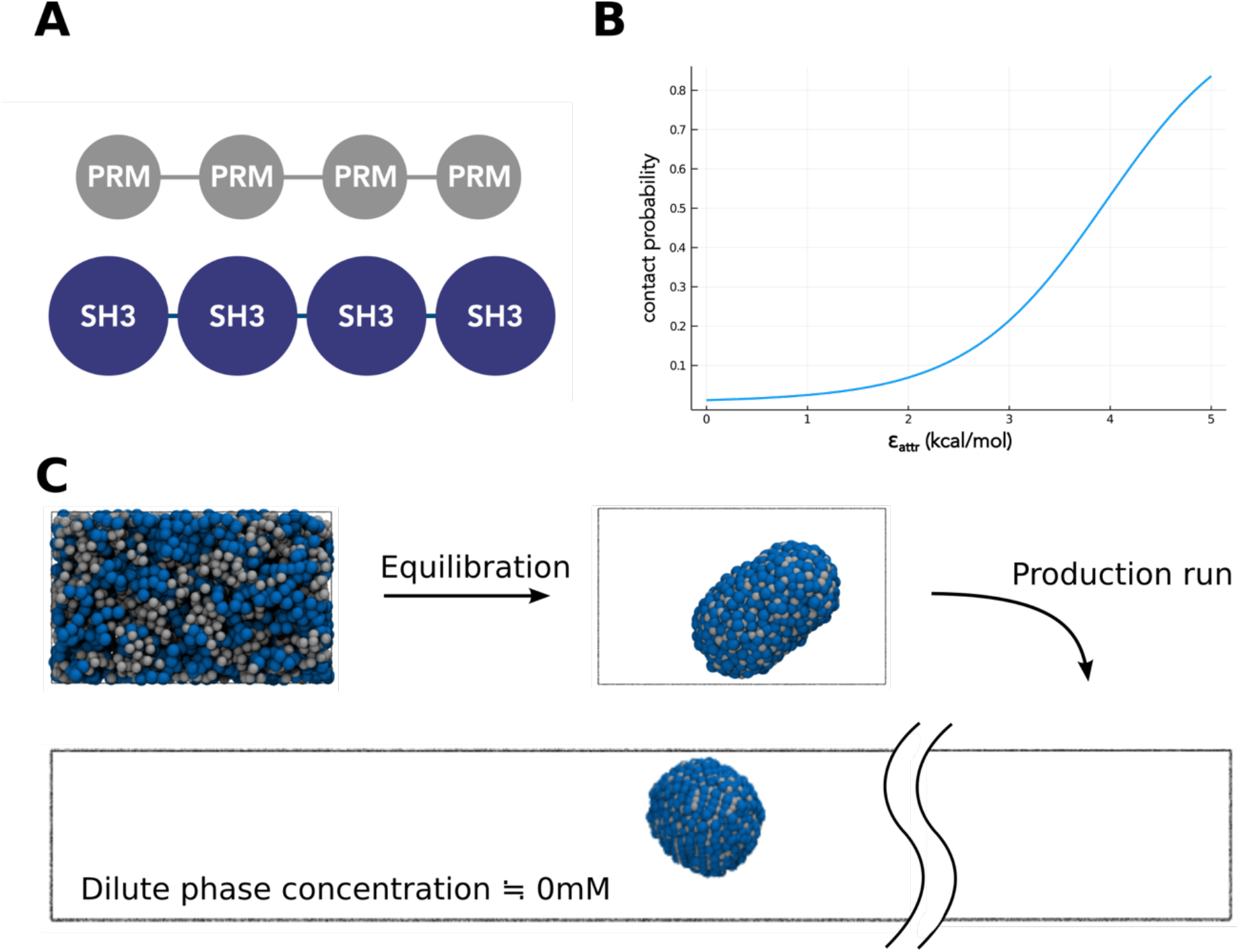
Pairwise interaction cannot reproduce appropriate dilute phase concentration. (A) A schematic representation of the SH3_4_-PRM_4_ system. SH3 domain and PRM motif are connected in tandem by flexible linkers. (B) The relation between ε_attr_ and the contact probability in 712 μM condition of SH3 and PRM. The 50% contact probability corresponds to the case *K*_*d*_ = 356 μM. (C) The snapshots of the SH3_4_-PRM_4_ system simulation with pairwise attractive interaction in the DC protocol. Blue particles represent SH3 domains and grey particles represent PRM domains.

In this set of simulations with a simple pairwise interaction, we found no SH3 or PRM molecules in the low-density phase, which had a density below the detection level (< 10 μM) for each type of molecule (Fig. 1C). This is in sharp contradiction to the critical concentration 100-200μM detected experimentally; in other words, the low-density phase in the phase-separated system should have a concentration of 100-200 μM.

We anticipated that this inconsistency could be because the pairwise interaction between SH3 and PRM excessively attracted interacting domains; in the simulation, one SH3 domain attracted several PRM domains and so did the PRM domain. When considering actual molecules, one PRM docks to one SH3, thus maintaining one-to-one stoichiometry. Ignoring stoichiometry may have led to over-compact interactions (we will show more direct evidence is shown below).

### Stoichiometric interaction can realize stoichiometry of protein-protein interaction

To solve the excessive attraction problem mentioned in the previous section, we propose a new non-pairwise interaction that explicitly considers stoichiometry (see Methods for the explicit equation). We call it the stoichiometric interaction (SI) potential. In the SI potential, the interaction strength for a specific particle is modulated by the number of interacting particles around that particle to avoid excessive stabilization under high-density conditions.

For example, consider the case in which one SH3 domain approaches two PRM domains. With the pairwise attractive interaction, the attraction is additive and the total stabilization is doubled (Fig. 2A, top). However, the SH3 domain strongly recognizes only one PRM in its binding site, whereas the second PRM can be attracted by weak nonspecific interactions on a different surface region of SH3. Therefore, the number of strong contacts should be one. As another example, we consider the case in which three SH3 domains are near by two PRM domians (Fig. 2A, bottom). With the pairwise attraction, there can be as many as six contacts that all additively stabilize the system. However, the actual number of strong contacts should be only two. Thus, the degree of stabilization must be approximately one-third of the estimate of the pairwise attraction. These two examples suggest that the strength of each contact can be adjusted to the appropriate strength by dividing it by the larger number of contacting particles of the two species for each pair of interactions. These provide reasonable estimates for most other cases of multiple contacts. To realize this, we propose stoichiometric interaction (SI) model, the explicit equation of which is given in the Methods section.

**FIGURE 2.**
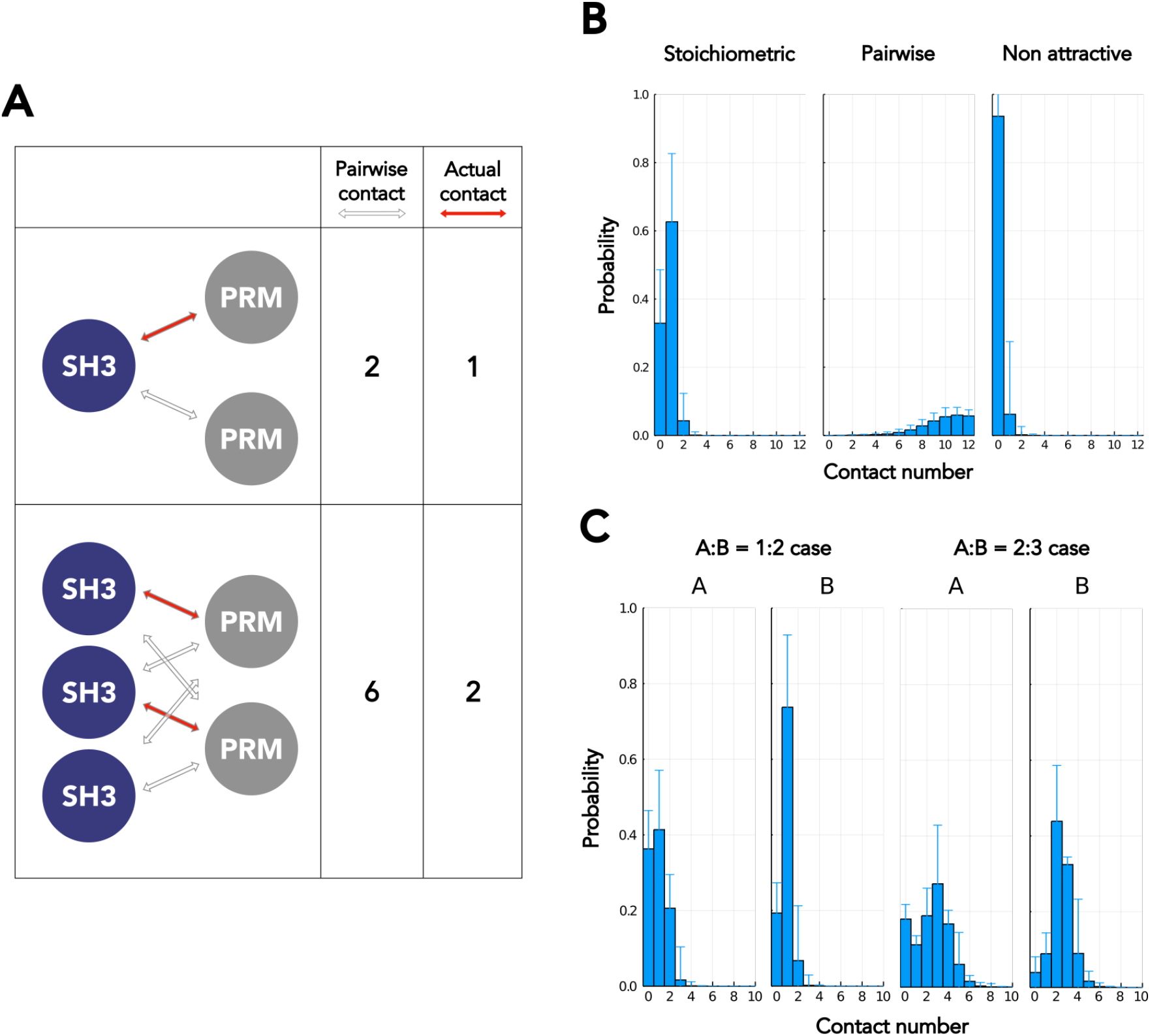
Stoichiometric interaction can repress over stabilization. (A) Schematic view of the pairwise attractive interaction and the one-to-one stoichiometric interaction. Upper panel: With one SH3 and two PRM’s in contacts, the pairwise interactions are doubled, but the number of actual one-to-one contacts is one. Lower panel: With three SH3’s and two PRM’s, there are six pairwise attractive interactions, whereas actual one-to-one contact are only two. (B) Distributions of the number of contacts per one SH3 (or PRM) in the stoichiometry assay for the SI model (left), the pairwise interaction (center), and with no attractive interaction (right). (C) Distribution of the number of contacts per A or B molecules in the case of non-1:1 valency. In this assay, the attractive interaction is only the SI model.

As a proof of concept, we performed simulations of a mixture of 1000 monomeric SH3 domains and 1000 monomeric PRMs in a cubic box with the SI model, as well as with the pairwise model as a control (Fig. 2B). We note that we tuned the interaction strength such that the dissociation constant *K*_*d*_ for the binary contact was identical between the two cases. For comparison, we also performed a simulation with no attractive interactions. In the case of the pairwise attractive interaction, a high-density state was excessively stabilized, and it became an aggregated state. The number of contacts per one monomer was distributed around 10 or higher (Fig. 2B, center). On the other hand, in the case of the SI model, the degree of contact was appropriately tuned, and the contact numbers per monomer became one or zero with high probability (Fig. 2B left).

The SI model can be extended to the cases of general valency that differ from one-to-one cases. We tested the case for A:B ratio of 1:2 and 2:3 (Fig. 2C). In the 1:2 stoichiometry case, the probability where the contact numbers per A particle were mostly between 0 and 2, and those per B particle were mostly 0 or 1, keeping the stoichiometric rule. For the 2:3 stoichiometry case, we found that the entire system tended to coalesce, which resulted in a broader distribution of the contact number with its peak at 3 for the A particles, and 2 for the B particles. Notably, once a large cluster was formed, even in the SI model, there are the cases where the number of particles that came close was greater than that designed in the SI model. However, the distribution also indicates that most of the contact numbers reflect the designed stoichiometry, suggesting that over stabilization was repressed by the SI potential.

### Both Stoichiometric interaction and pairwise interaction are needed to realize appropriate concentration of the dilute phase

In the previous section, we confirmed that the SI model could realize the desired stoichiometric interactions in test cases. Then, we returned to our target system of the binary mixture of SH3_4_ and PRM_4_, performing simulations like those mentioned above using the SI model. We set the SI parameter such that the domain-domain dissociation constant became *K*_*d*_ = 356µM as in the previous case. The simulation resulted in a phase-separated equilibrium in which the low-density phase had a concentration of approximately 2000 μM (Fig. 3D, bottom). This concentration in the low-density phase was an order of magnitude larger than the experimentally observed threshold concentration of 100-200 μM (39). The SI model limits the density of the interaction network, resulting in moderate transition into the phase-separated system. We note that the purely pairwise attractive interaction resulted in excessive clustering into an aggregate with no detectable concentration in the low-density phase (Fig. 3D, top), whereas the pure SI model resulted in a weak phase separation with too high a concentration in the low-density phase (Fig. 3D bottom). In fact, the real molecular system should have, both, specific 1:1 interaction, by which SH3 recognizes PRM, and non-specific weak and promiscuous interactions.

**FIGURE 3.**
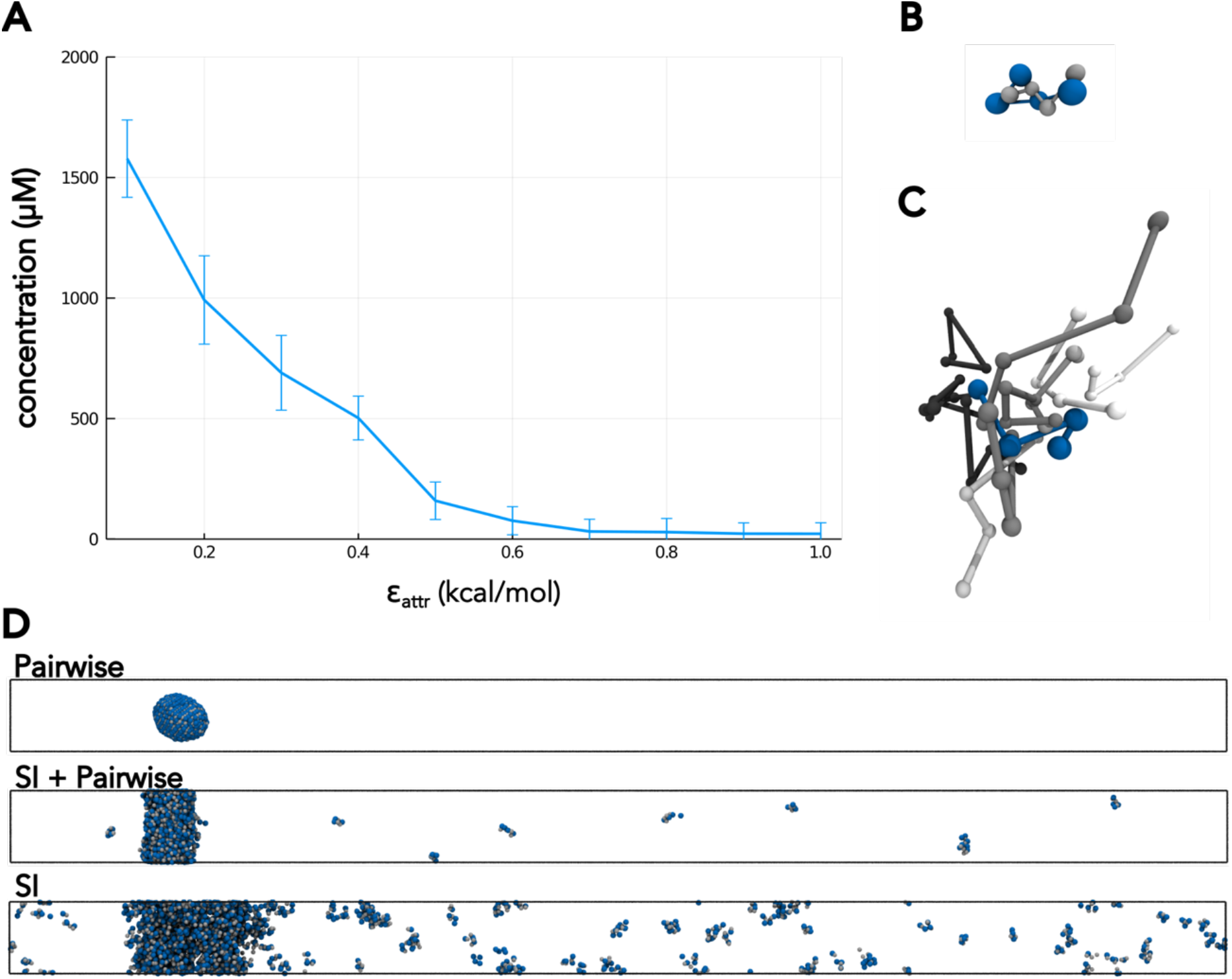
The system with both Stoichiometric and pairwise interaction can reproduce appropriate dilute phase concentration of SH3_4_ PRM_4_ mixed system. (A) The strength of pairwise attractive interaction vs critical concentration. SI interaction strength corresponds to Kd=356 μM. Error bar shows standard deviation. (B) The snapshot of a part of the dilute phase molecules. One of SH3_4_ molecules and PRM_4_ molecules around that SH3_4_ are visualized. (C) The snapshot of a part of the condensed phase molecules. One of SH3_4_ molecules and PRM_4_ molecules around that SH3_4_ are visualized. PRM_4_ molecules in different colors have different nearest domains of SH3_4_ molecules. (D) The snapshot of three types of systems. In the top system, only pairwise interactions are included. Middle system shows inclusion of both pairwise interactions and SI (Movie S1). Bottom system depicts a system with only SI. Blue spheres represent SH3 domains, and grey spheres represent the PRM motives. The linker between each domain and motif is not displayed for visibility.

To this end, pairwise interactions were added to the SI model. The strength of this pairwise interaction ε_attr_ was tuned such that the concentration of the low-density phase matched the value observed in the experiment. Finally, we ended up with ε_attr_ = 0.5 (kcal/mol) (Fig. 3A). With this balance, we observed that the phase-separated system was stabilized with the concentration of the low-density phase being 159 µM (Fig. 3D, middle, Movie S1).

Movie S1. MD simulation of the LLPS of the SH3_4_ and PRM_4_ with the SI and pairwise interactions (corresponding to the middle panel of Fig. 3D).

### The phase diagram of phase separation

With the tuned parametrization of the SI and pairwise interactions, we obtained the phase diagram of the phase separation for the SH3_4_-PRM_4_ systems. In this set of simulations, we prepared in-total 11 average concentration ratios of SH3_4_ and PRM_4_, ranging from 40% to 60% SH3. When the ratio of the SH3 domain was over 50%, the critical concentration, which corresponds to dilute phase concentration, for the PRM domain was approximately 200 μM; the critical concentration of SH3 increased as the SH3 average concentration increased (Fig. 4A). On the other hand, when the ratio of the PRM motif was over 50%, an opposite tendency was observed; the critical concentration of the SH3 was approximately 200 μ M, while the critical concentration of PRM increased as the average concentration of PRM increased. We obtained the phase diagram (Fig. 4A). The phase diagram and the critical concentrations matched the results of the experiment (39). Representative snapshots are shown in Fig. 4C.

**FIGURE 4.**
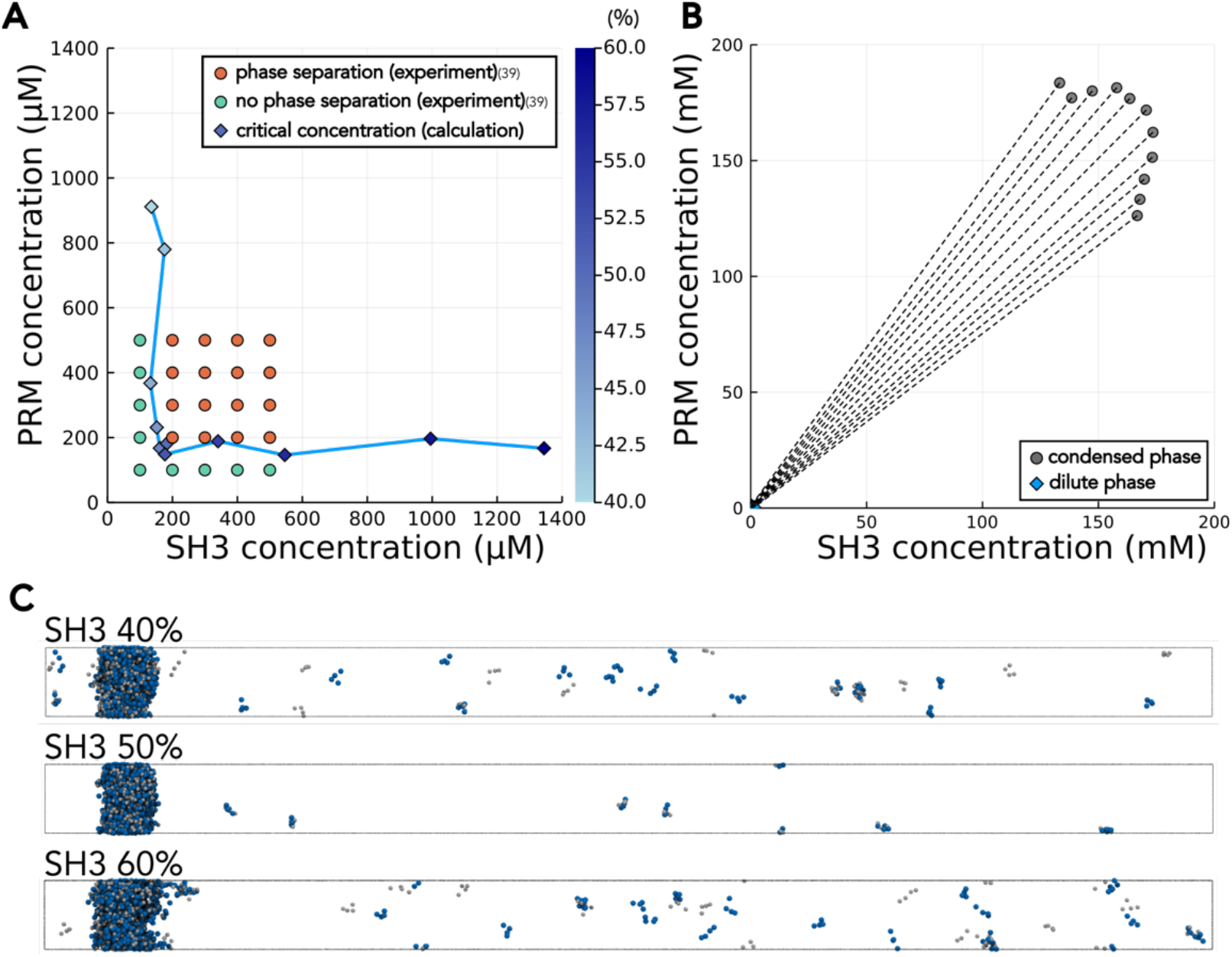
The system with both stoichiometric and pairwise interaction can reproduce phase diagram of SH3_4_ PRM_4_ mixed system. (A)The phase diagram for the phase-separation of the SH3_4_-PRM_4_ system using experiments and simulations. Diamonds are critical concentrations in the simulation. The color of the diamonds indicates the ratio of SH3 and PRM domains. Circles show the result obtained from previously reported experiments (39); Orange indicates phase separation, while green indicates the no phase separation. (B)Tie lines (dashed lines) of simulated SH3_4_-PRM_4_ systems. The dilute phase concentrations same as those in diamond points in (A) all aggregated to the bottom left points near the origin. Circles represent the condensed phase concentrations. (C)Snapshots of the cases with the ratios of SH3 domains being 40% (top), 50% (center), and 60% (bottom). Blue particles indicate SH3 domains and grey particles indicate PRM motives. The linker between domains is not shown for visibility.

Recent studies suggested that complex molecular interactions of multi-component systems can be reflected by the tie lines (8, 41), and thus we drew them (Fig. 4B). A tie line can be drawn by connecting concentrations of the dilute phase and those of the condensed phase in a phase-separated simulation. We found that the tie lines diverge from dilute phases to condensed phases. This is consistent with the observation in a recent work of Chan’s group (42). In their work, they clarified that the two-component system containing stochiometric complex and promiscuous interaction exhibits the divergent pattern of tie lines. We also found that the system with only the stoichiometric attractive interaction leads to the divergent tie line pattern (Fig. S2), which is distinct from the previous study (42).

### Liquid dynamics in the phase-separated system

Thus far, we have shown the characterization of the equilibrium phase behaviors of the SH3_4_-PRM_4_ systems with the SI potential augmented by the pairwise interaction. Next, we address the dynamics of the same system. The diffusion coefficients of the SH3 domain and PRM in the dilute and condensed phases were calculated from the mean square deviation (MSD) plots (Fig. 5A). To obtain the MSD, we first divided each trajectory into 10^5^ step windows without overlap, chose a representative particle from each of the four categories, namely SH3 and PRM in the dilute and condensed phases, and calculated the MSD. With the SI model augmented by pairwise term, all four MSD curves with respect to the MD time step showed approximately linear dependence with non-zero slopes (Fig. 5A, S2), suggesting the fluidity of both the dilute and condensed phases. Subsequently, MSD curves were fitted using linear regression. The estimated diffusion coefficients for the SH3 domain were (1/6) × 2.56 Å^2^/ MD-step in the dilute phase and (1/6) × 2.91 × 10^− 1^ Å^2^/ MD-step in the condensed phase. For the PRM, the estimated diffusion coefficients were (1/6) × 2.57 Å^2^/ MD-step in the dilute phase and (1/6) × 3.39 × 10^− 1^ Å^2^/ MDstep in the condensed phase (Fig. S3). Thus, the estimated diffusion coefficients in the dilute phase was one order of magnitude larger than those in the condensed phase, which was qualitatively in agreement with the experimental results (39), although quantitatively, the difference was smaller in the simulations.

**FIGURE 5.**
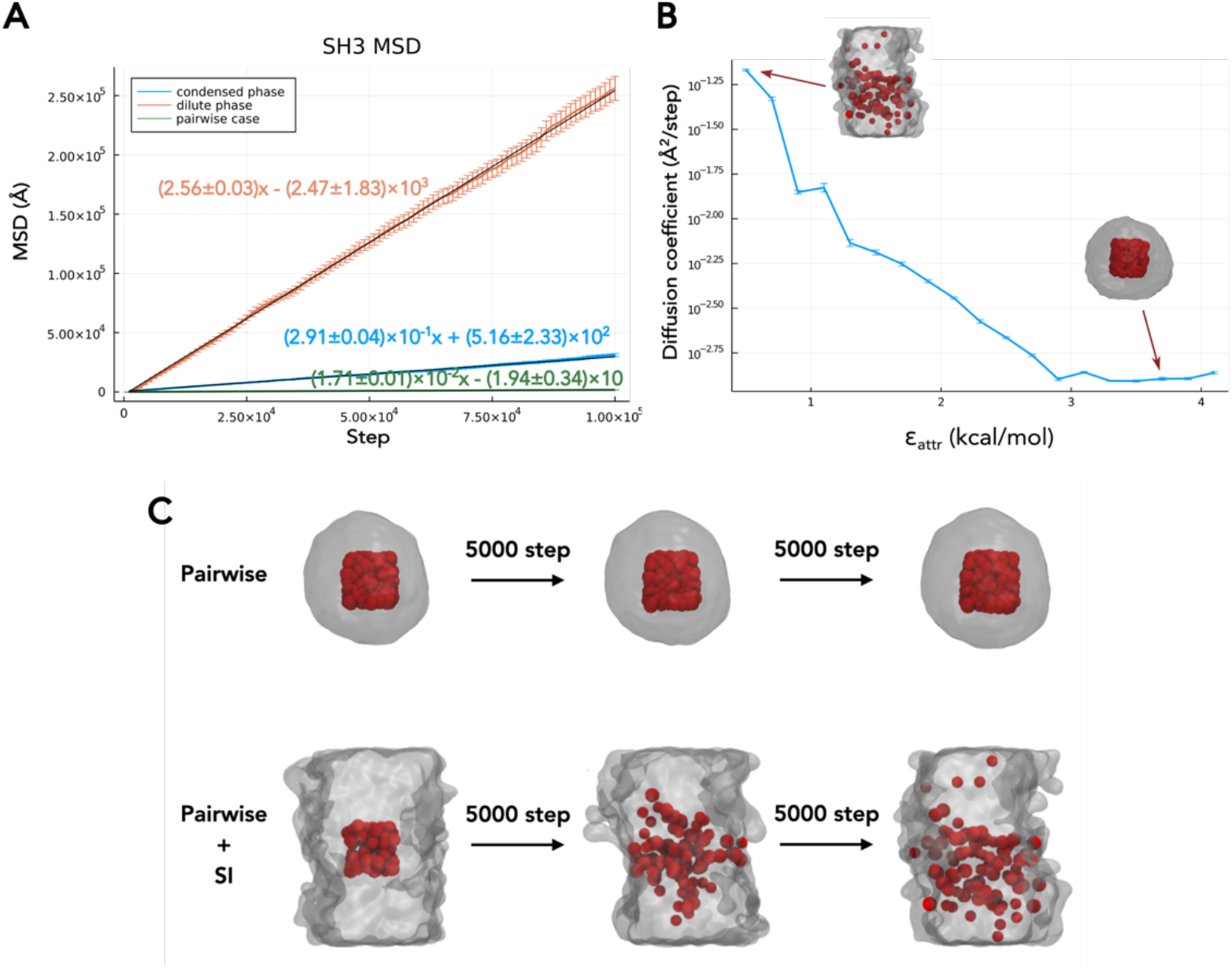
The system with both stoichiometric and pairwise interaction shows the liquid activity of condensed phase. (A) The MSD curves of SH3 molecule with respect to the MD step. The asymptotic regions of the curves were fitted via the linear regression. Orange and blue indicate the dilute and condensed phases with the balanced SI and pairwise interactions, respectively. Green indicates the condensed phase with only pairwise interactions. (B) The diffusion coefficient of SH3 molecule along the strength of the pairwise attractive interaction. The snapshots in the graph are of corresponding FRAP-like simulation, tracked 10,000 MD step cases. (C) *In silico* FRAP-like simulation (Movie S2 and S3). Grey area shows the condensed phase. At a time, all the particles in a cubic box are labeled. These particles were tracked after 5,000 MD step and 10,000 steps. Top, with only the pairwise interaction; bottom, with the balanced SI and pairwise interaction.

For comparison, we also estimated the MSD of the same system simulated solely using pairwise attractive interactions (Fig. 5A, S2, green). The MSD of the SH3 (or PRM) increased much more slowly with MD time than in above case. We note that, in this case, all the molecules were assembled into one aggregate (cluster) and that the aggregate itself could diffuse.

To address whether this difference in MSD slopes comes from qualitatively different dynamics, such as the grass-transition, or not, we performed simulations with varying ratios of SI and pairwise attractive interactions (Fig. 5B). In these simulations, while the sum of *ε*_*SI*_ and *ε*_*attr*_ was kept the same as that of the above simulations, their ratio *ε*_*attr*_/*ε*_*SI*_ was modulated. We found that, as the ratio *ε*_*attr*_/*ε*_*SI*_ increases, the diffusion coefficient decreases (Fig. 5B). The slope in the semi-log plot is nearly constant from *ε*_*attr*_ = 0.5 (i.e., the above default case) to 2.9, whereas the diffusion coefficient becomes nearly constant above *ε*_*attr*_ = 2.9. The singular behavior at the critical value *ε*_*attr*_ = 2.9 implies the phase transition; liquid-like dynamics below that point, while glassy dynamics above the threshold.

To distinguish the diffusion within the cluster from that of the entire cluster, we subsequently performed an *in silico* FRAP-like analysis (Fig. 5C, Movie S2 and S3). In a well-equilibrated system, we first set a cubic box at the center of the formed cluster and labeled all particles in the box at a time (Fig. 5C, left). We then tracked all labeled particles at the subsequent time. We found that the simulation with only pairwise attractive interactions did not exhibit noticeable diffusion during 10,000 MD steps (Fig. 5C, top). In contrast, the simulation with balanced SI and pairwise interactions showed rapid diffusion of the labeled particles over the 10,000 MD steps (Fig. 5C, bottom). These results clearly showed that the balanced SI and pairwise interactions generated a liquid droplet, while the purely pairwise interactions did not exhibit fluidity.

Movie S2. *In silico* FRAP-like simulation with only pairwise interactions (corresponding to the top panel of Fig. 5C).

Movie S3. *In silico* FRAP-like simulation with the SI and pairwise interactions (corresponding to the bottom panel of Fig. 5C).

### Spontaneous formation of the phase-separated droplet

Next, we investigated the spontaneous formation of droplets from a uniformly distributed configuration of the SH3_4_-PRM_4_ systems with balanced SI and pairwise interactions (Fig. 6A, Movie S4). In this simulation, the DC method was not used. Instead, we set up a box filled with 375 randomly distributed SH3_4_ and PRM_4_ molecules and started the simulation (Fig. 6B, top). The total step of simulation consisted of 5.0×10^7^ MD steps. Within the initial 10^5^ MD steps, nearby molecules quickly formed small clusters (Fig. 6B, second panel). These small clusters subsequently collided with each other and coalesced into larger clusters. At the 10^6^ MD step, only there large clusters remained together with a few molecules in the dilute phase (Fig. 6B, third panel). At the 10^7^ MD step, only two clusters remained with the largest cluster size being ∼600 molecules. At around the 1.6×10^7^ MD step, all the clusters merged into one droplet, the high-density phase containing ∼700 molecules, which is the converged size of the droplet (Fig. 6A). Since this point, we observed occasional dissociation of a pair of molecules from the high-density phase and association of a pair of molecules from the low-density phase to the high-density phase.

**FIGURE 6.**
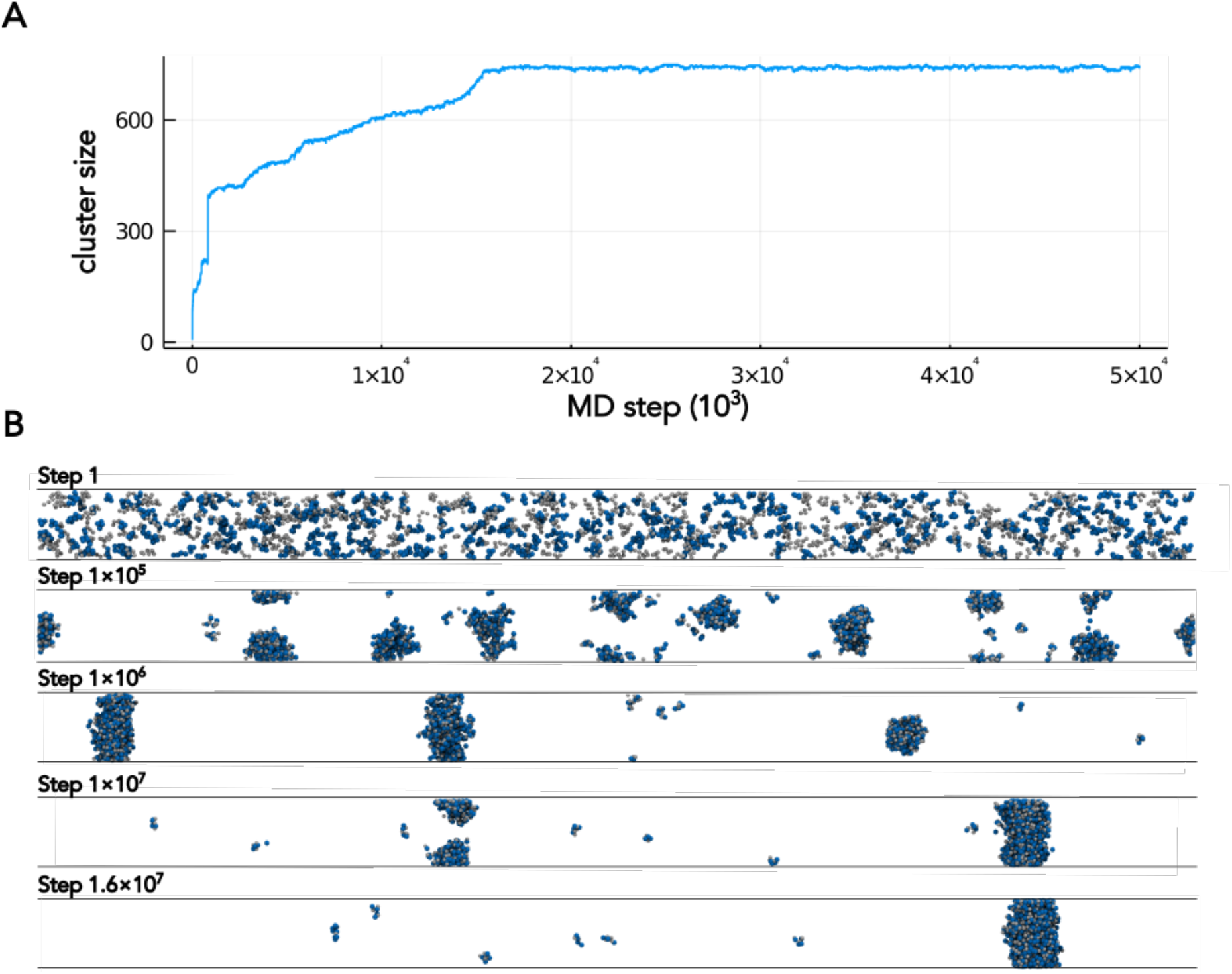
The simulation from the configuration with uniformly distributed molecules to equilibrated phase-separated system. (A) Time series of the maximum cluster size of the simulation start from a uniform distribution. Cluster size represents the number of SH3 and PRM molecules in the cluster. (B) Snapshots of the trajectory start from a uniform distribution (Movie S4). Blue particles represent SH3 domain and grey particles represent PRM motives. The linker between domains is not shown for visibility.

Movie S4. Spontaneous phase separation of the SH3_4_ and PRM_4_ with the SI and pairwise interactions (corresponding to Fig. 6).

### Dependence on the valency

It has been reported that phase behavior depends on the valency of the interacting molecules (39). For the SH3 and PRM systems, the threshold for the phase separation was investigated for the cases of variable valency, such as SH3_3_-PRM_3_, SH3_3_-PRM_4_, SH3_3_-PRM_5_ etc. We examined whether the balanced SI and pairwise interaction model could reproduce this result. We performed DC simulations and analyzed the phase diagrams for various combinations of valency, i.e., SH3_n_-PRM_m_ with n and m being 3, 4, or 5 (Fig. 7).

**FIGURE 7.**
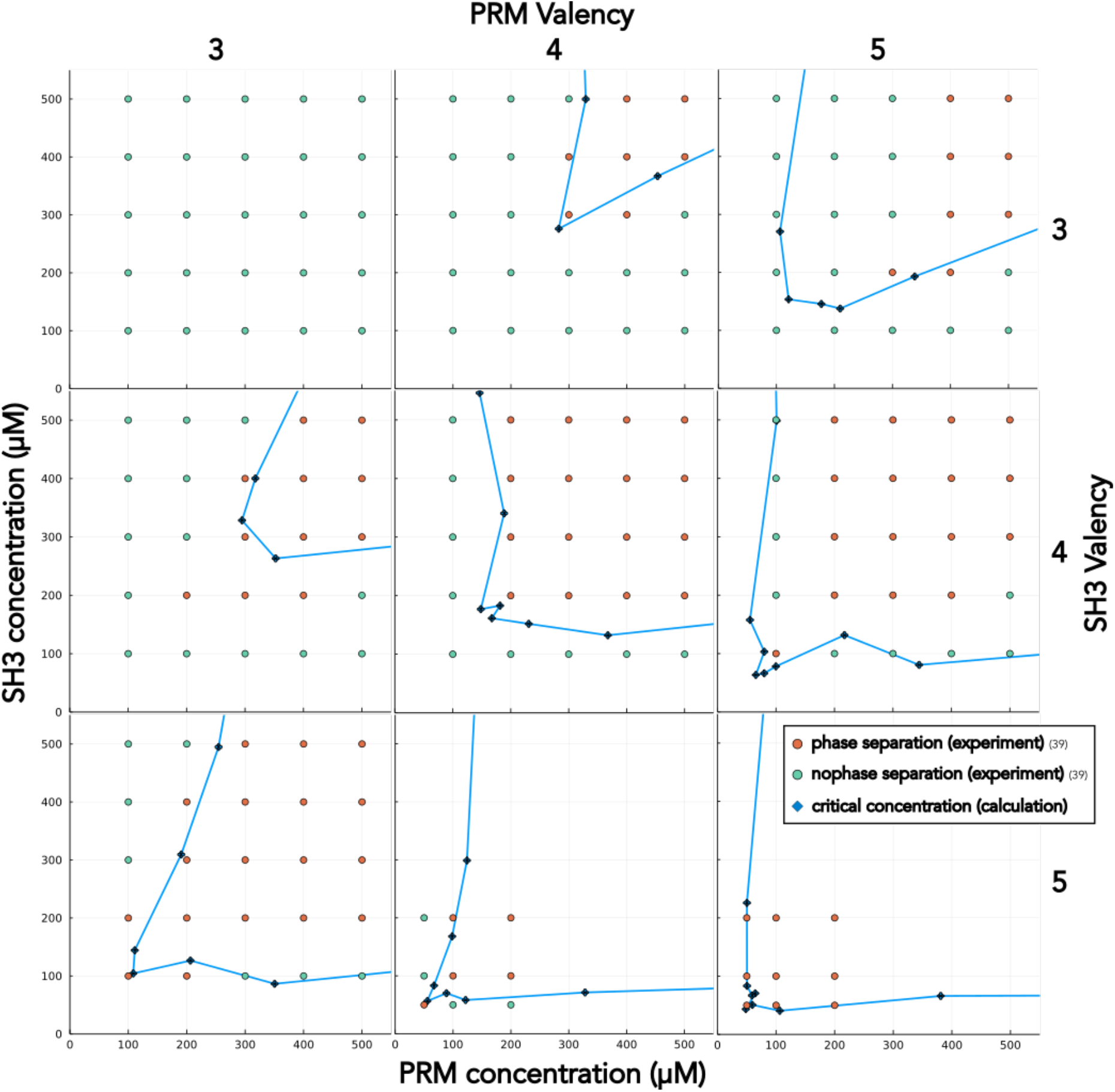
Phase diagrams depicting phase separation for various valences of SH3 and PRM. The valency of SH3 is 3, 4, and 5, for the top, middle, and bottom rows. The valency of PRM is 3, 4, and 5 for the left, middle, and right columns. In each panel, the vertical axis of each graph represents the SH3 domain concentration, and the horizontal axis represents the PRM concentration. Blue diamonds represent critical concentrations in the simulation. Circles show the results obtained from previously reported experiments (39); Orange, the phase separated; green, the uniform phase.

For the case where the valency of SH3 and PRM was the same, i.e., n=m, the threshold concentration for the phase separation for the same concentration of SH3 and PRM module case decreased as the valency increased from 3 to 5. The thresholds were ∼1117 µM (SH3) and ∼1094 μM (PRM) for n=3, ∼183 μM (SH3) and ∼182 μM(PRM) for n=4, and ∼70 μM (SH3) and ∼64 μM (PRM) for n=5. When the valency of SH3 was smaller than that of PRM, a higher concentration of PRM was required to induce the phase separation for cases with low SH3 concentrations. However, when the valency of PRM was smaller than that of SH3, the opposite trend was observed.

## CONCLUSIONS

Liquid-liquid phase separation (LLPS) forms biomolecular condensates; the fluidity and complexity of their components makes molecular simulation approach indispensable. In the simulation approach, to overcome the problem of the mesoscopic model for biomolecular condensates composed of multi-domain proteins, we have proposed the stoichiometric interaction (SI) potential that explicitly considers the valency of the interacting domains in the mesoscopic model. First, we examined whether the SI potential maintains the valency for the test system of binary molecular solutions, and found that the SI potential, but not the ordinary pairwise interaction potential, can regulate the valency well. For the n:m stoichiometry case, this stoichiometry was reproduced in the simulation using the SI potential.

Then, we used the SI potential for simulations of a binary mixture of tandem repeats of the SH3 domain and PRM; the phase behavior for this system has been studied well experimentally. We found that to reproduce the LLPS phase behavior quantitatively, we had to add a pairwise interaction potential to the SI potential. In this mixed potential, pairwise interactions can be interpreted as promiscuous interactions that do not have a well-defined valency. This result suggests that the promiscuous interactions, in addition to the stoichiometric one-to-one interactions, are indispensable for LLPS in the SH3-PRM system. For domain-domain stoichiometric interaction systems, the contribution of promiscuous interactions was also reported in a previous theoretical research of Chan’s group (42). This together strongly suggests important role of the promiscuous interaction even in LLPS of domain-domain interacting systems.

By estimating the diffusion coefficient and using an *in-silico* FRAP-like simulation, we confirmed the fluidity of the condensed phase realized by the mixed SI and pairwise potential and the non-fluidity of the case with the pairwise potential alone. Moreover, we examined the case of different numbers of interaction domains, including SH3_3_-PRM_3_, SH3_3_-PRM_4_, …, SH3_5_-PRM_5_, and found that we could reproduce the phase diagrams for all but one case.

## AUTHOR CONTRIBUTIONS

YM and ST conceived the study. YM performed the simulations and analyzed the data. TN supported software development. YM and ST wrote the manuscript. All the authors have read and checked the manuscript.

## ACKNOWLEDGMENTS

The study was supported partly by JSPS KAKENHI grants 19J23454 (YM), 20H05934 (ST) and 21H02441 (ST), and partly by JPMXP1020200101 as “Program for Promoting Researches on the Supercomputer Fugaku” (ST).

## SUPPORTING INFORMATION

### Implementation of Stoichiometric Interaction Potential

A naive implementation of the SI potential can make the calculation impractical. We can avoid it. The derivative of the SI potential with respect to the coordinate *r*_*k*_ is

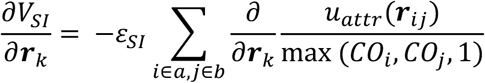

The derivative becomes, if *k* ∈ *a* case,

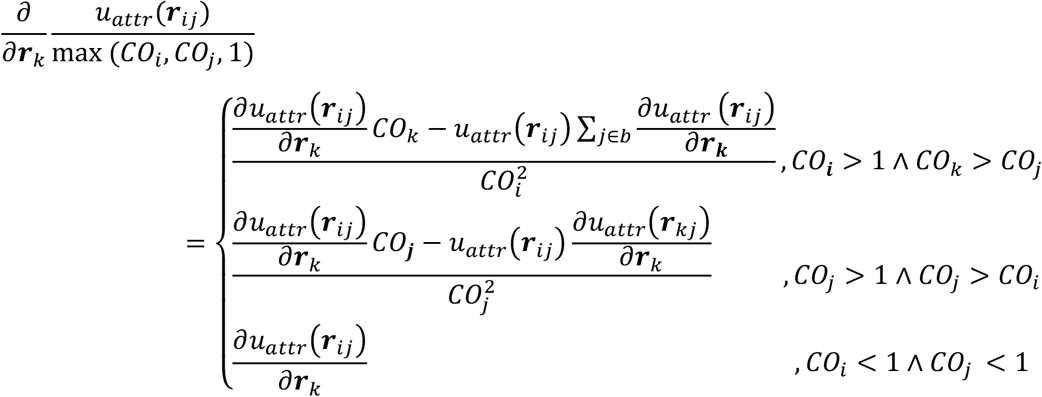

and if *k* ∈ *b* case,

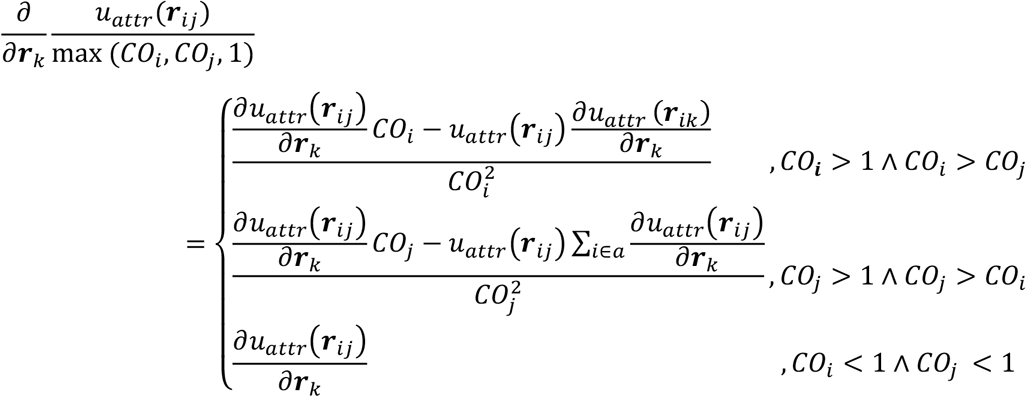

All derivatives are composed of *u*_*attr*_ (*r*_*ij*_) and 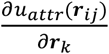. Once we have all *u*_*attr*_ (*r*_*ij*_) and 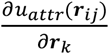 values, we can calculate the derivative of the SI potential by *O*(*n*) (n is the number of particles participant to SI). The order of *u*_*attr*_ (*r*_*ij*_) and 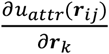 calculations correspond to pairwise interaction, so the order of all calculations is also the same as that. We need buffering *u*_*attr*_ (*r*_*ij*_) and 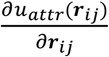 before 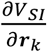 calculation. Since these are sparse matrices, we stored these value using Compressed Sparse Row scheme. We also use Cell List for the calculation of buffering values.

### Supplemental figures

**Figure S1.**
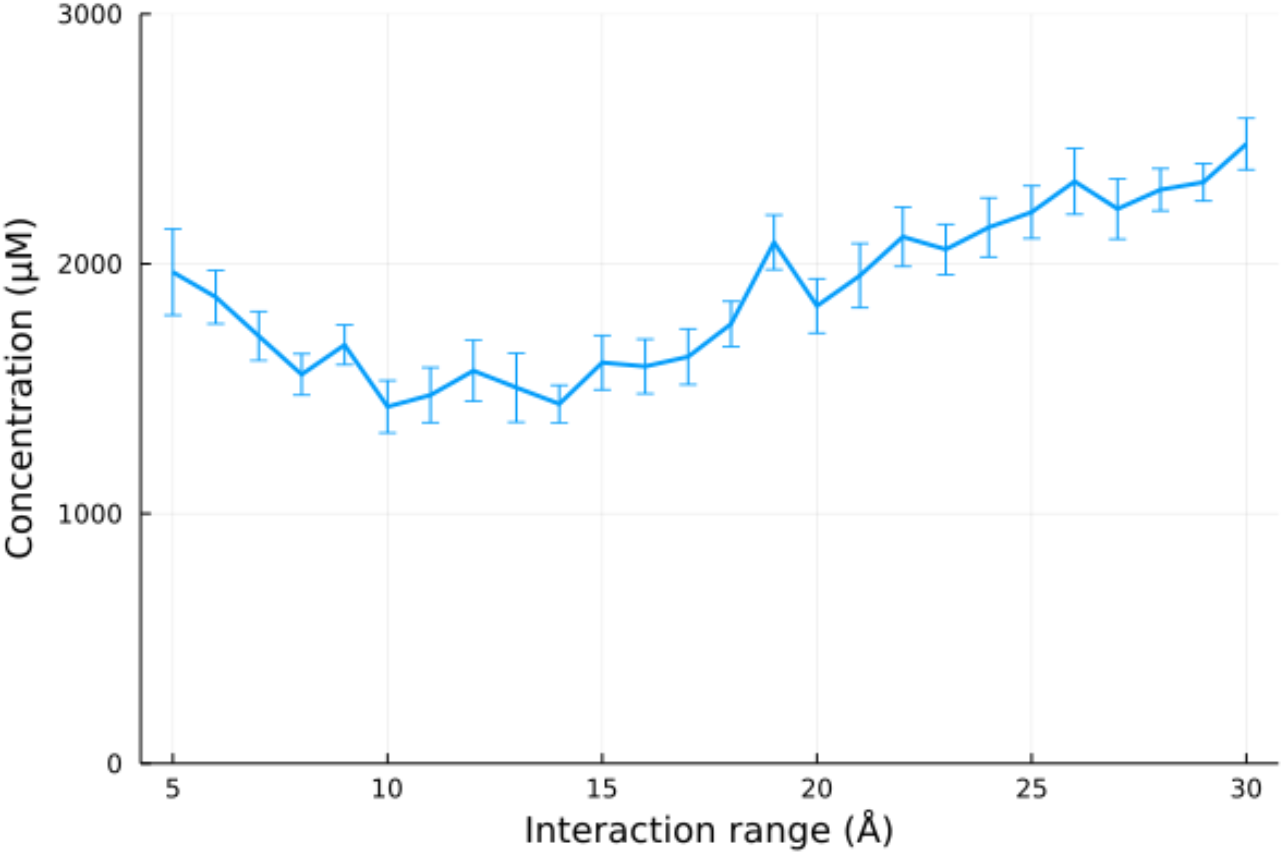
The dependence of the LLPS on the attractive interaction range in the SI potential. The graph is to address the dependence of the LLPS behavior on the attractive interaction range of the SI potential. The simulation was conducted by the DC method. The attractive interaction was only the SI potential. ε_SI_ was set as 3.91 (kcal/mol). The horizontal axis shows the attractive rangeΔr in u_attr_() in the SI potential. The vertical axis shows the dilute phase concentration in the LLPS. The error bars show the standard error. The result shows that the results are insensitive to the value of the attractive interaction range.

**Figure S2.**
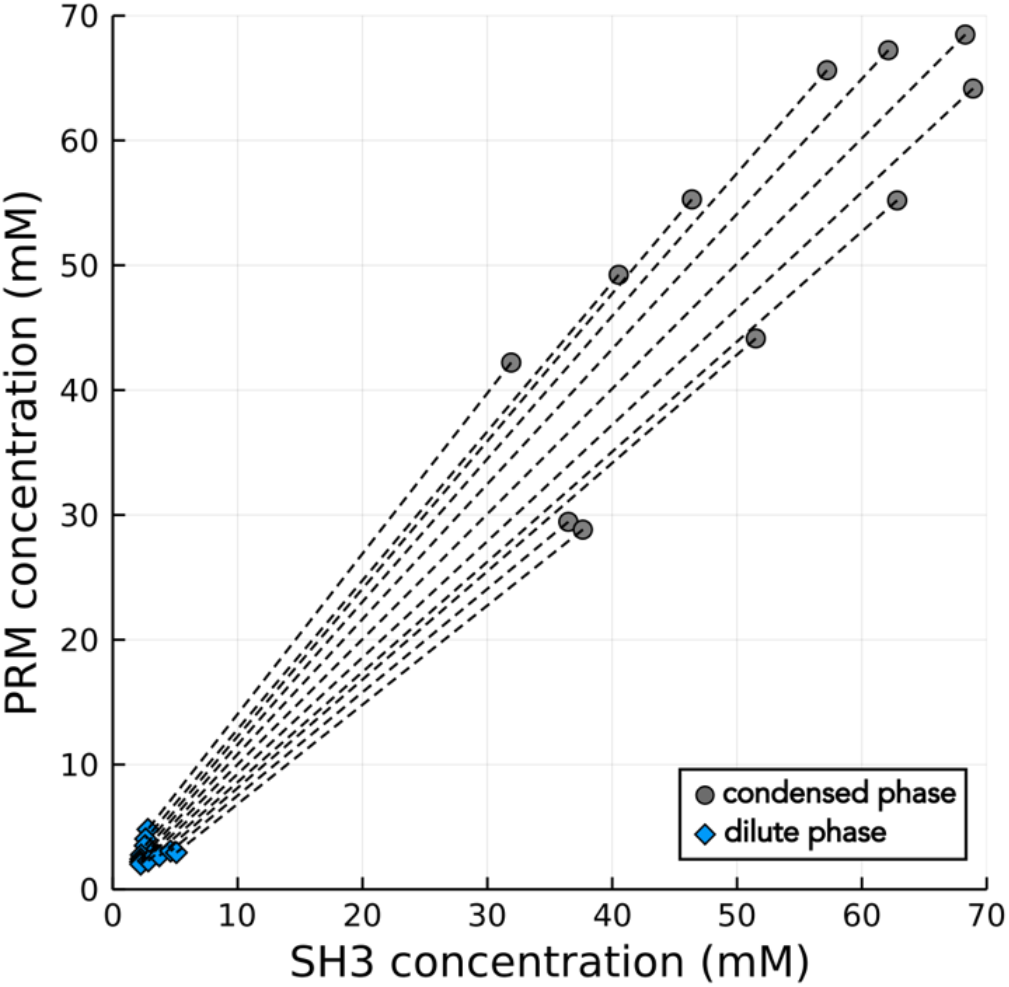
The tie lines of simulated SH3_4_-PRM_4_ system with only the SI potential. Each tile line connects the concentration of the dilute phase (diamond) and that of the condensed phase (circle). Different tie lines correspond to different ratios of SH3 domain to PRM concentrations, every 4% from 40 to 60%.

**Figure S3.**
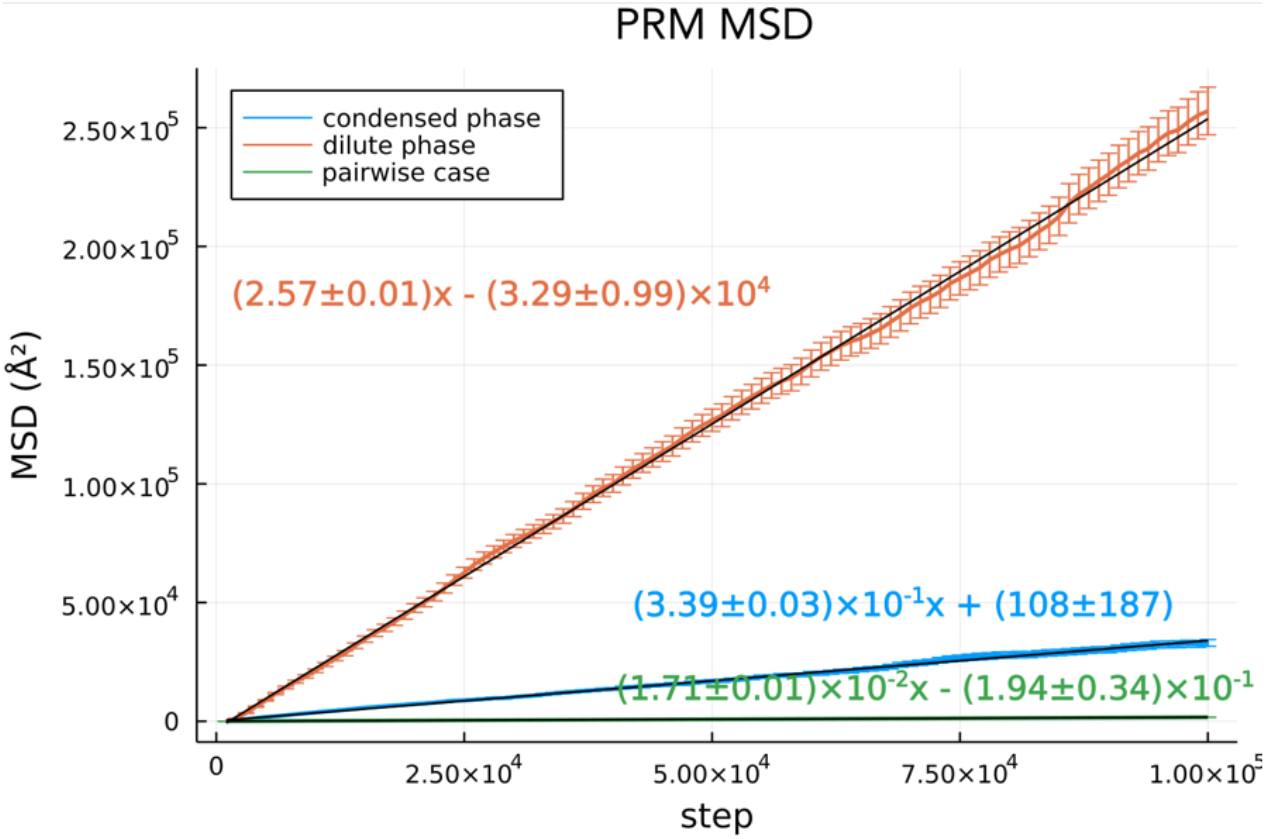
The MSD curves of PRM molecule with respect to the MD step. The asymptotic regions of the curves were fitted via linear regression. Orange and blue indicate the dilute and condensed phases with the balanced SI and pairwise interactions, respectively. Green indicates the condensed phase with only pairwise interactions. The error bars show the standard error.

